# Ubigen: Interactive Ranking and Enrichment Test for Ubiquitously Expressed Genes

**DOI:** 10.1101/2022.07.28.501880

**Authors:** Elias Projahn, Steffen Möller

## Abstract

We provide an interactive web service to score and rank human genes by their abundance and invariance of expression across all samples in the GTEx database. Users may provide a list of genes to have these scored and ranked accordingly. A one-sided Wilcoxon rank sum test determines a p-value to indicate if the genes from the list score higher than expected by chance. It is possible to interactively control the parameters and instantly observe the effects on the ranking for each of the submitted genes of interest. A genome-wide sliding-window analysis finds that the genes with both the highest and the lowest ranks feature more GO term annotations, indicating that the scoring carries biological relevance.

**Availability and Implementation:** https://ubigen.uni-rostock.de

---

An open question since the advent of high-throughput transcriptomics has been the identification of housekeeping genes^1^. In the literature, the terms housekeeping, constitutive and ubiquitous^2^ are often used interchangeably, loosely differentiating genes based on their abundance, using expression data as the only source of information. Ubiquity of expression does not necessarily have a biological implication such as a function in cellular “housekeeping” or a specific mode of gene expression control. Instead, it is based on a statistical definition: Genes are called ubiquitous if they are expressed at a sufficiently high level in a sufficient fraction of samples and optionally, if they are expressed at a high level and/or with low variation *across tissues and conditions.* Thus, ubiquitous genes are also of interest for quality-checking of expression data.

Over the last decade, several studies used large expression datasets to identify “housekeeping” genes. A common approach has been to filter for genes that are expressed above a certain threshold throughout all available samples, irrespective of variation^3,4^. In other studies, additional constraints on the variation of expression were added^5,6^. In order to incorporate the means to compare different ways of approaching the concept of ubiquity, we have developed a web interface that interactively integrates multiple scoring functions to allow representing both concepts (Table 1).

**Table 1:** Selection criteria for ubiquitous genes. Genes are ranked by a ubiquity score that is the weighted sum from multiple criteria. The user may adjust the weights. By default, the fraction of samples featuring high expression of the gene and the two secondary criteria are considered as two blocks that contribute equally to the score. Within the second block, the mean expression is weighted positively and the standard deviation is weighted negatively.

| Criterion | Description | Default Weighting |
| --- | --- | --- |
| Fraction of samples with high expression | Fraction of samples that express the gene based on a specified threshold. The threshold is computed separately for each sample and corresponds to the 95<sup>th</sup> percentile of expression within that sample. It is possible to select the median (i.e., the 50<sup>th</sup> percentile) or zero as reference values, too. | 50% |
| Mean expression | A mean expression level (logarithmic) is determined for each gene across all samples. | 25% |
| Standard deviation | Analogous to the mean expression, the standard deviation is determined for each gene across all samples. | -25% |

Our analysis is based on expression data from the GTEx project^7^ which, at the time of writing, encompasses 17,382 samples from different human tissues. A primary criterion for the ranking of genes is the *fraction of samples* in which an individual gene is highly expressed. By default, we define “highly expressed” as an expression above the 95^th^ percentile within a sample. Alternatively, it is offered to qualify “highly expressed” genes based on the median expression level per sample or based on whether they are expressed at all. In addition to that, it is possible to prefer genes that have a high mean expression and/or low standard deviation across all samples. These different criteria (Table 1) are combined into a single ubiquity score for each gene by a weighted average.

This project provides a simple interface that facilitates the selection of ubiquitous genes and enables researchers to rank their genes of interest accordingly (Figure 1). This provides an almost immediate^8^ visualization of their ubiquity and the derivation of a p-value (Wilcoxon rank sum test) with the alternative hypothesis that the custom genes have higher scores than all other genes. It also integrates the presentation of enrichment analyses with g:Profiler^9^.

**Figure 1:**
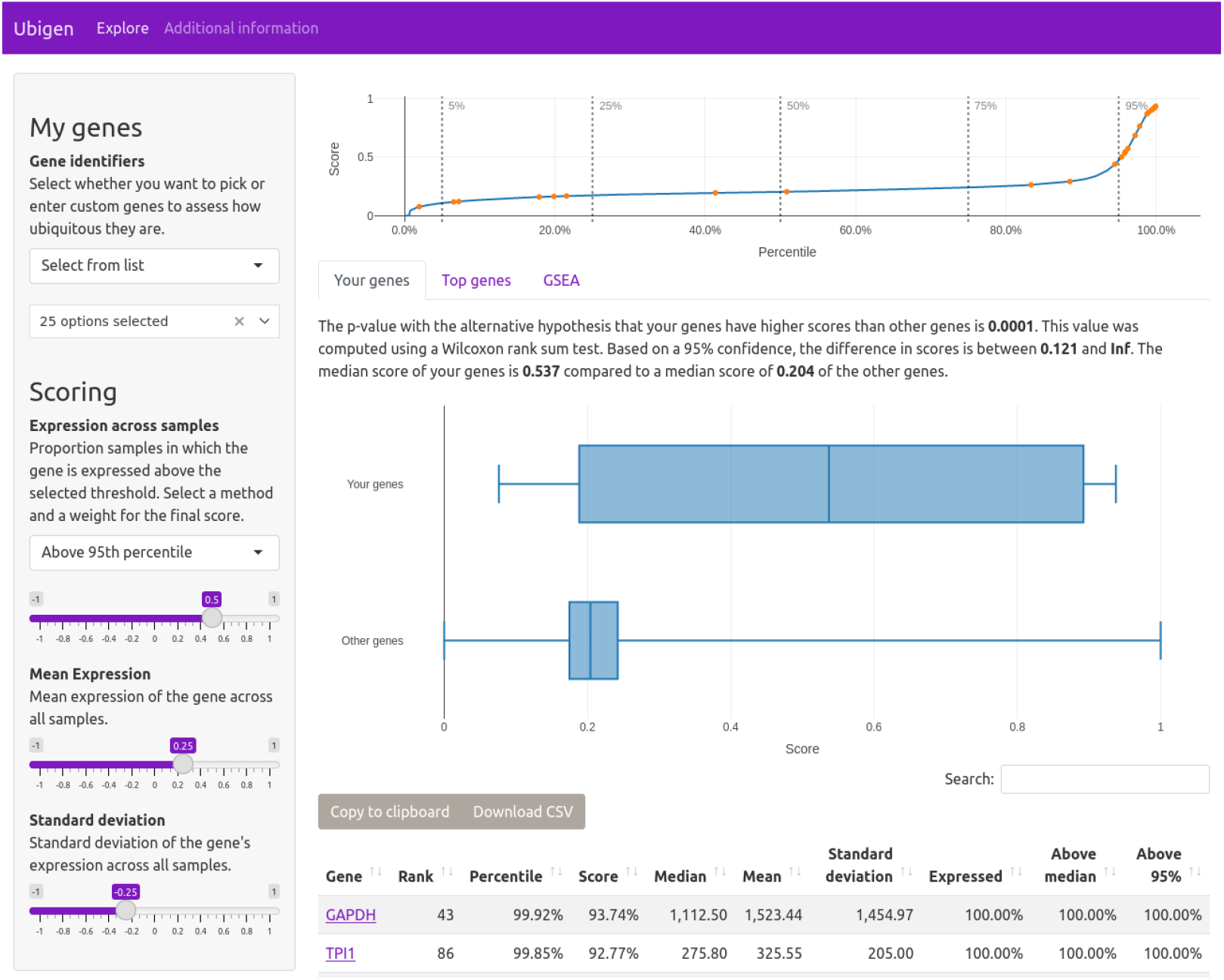
Screenshot of interactive web user interface. Parameters of the ranking can be adjusted instantaneously to specify how to compute the genes’ ubiquity score, thus enabling a user-controlled weighting of the different criteria (Table 1). Multiple interactive plots are included to visualize the ranking. A prominent feature is the inclusion of custom genes to assess their possible categorization as ubiquitously expressed genes. In the pictured case, all genes involved in glycolysis according to the KEGG pathways database [KEGG:`hsa00010+M00001`] were selected as an example.

Our initial hunch was that genes that are ubiquitous may be difficult to characterize in terms of the biological processes they are involved in. However, we found that the most ubiquitous genes are enriched for a large number of GeneOntology terms. Figure 2 presents the number of associated terms in a non-overlapping sliding window of 1000 genes along the ranking of ubiquity with default parameters. We see that the genes at either end of the spectrum are exceptional, which also supports the validity of this analysis. The web interface interactively presents an extension of Figure 2 for all data sources covered by g:Profiler.

**Figure 2:**
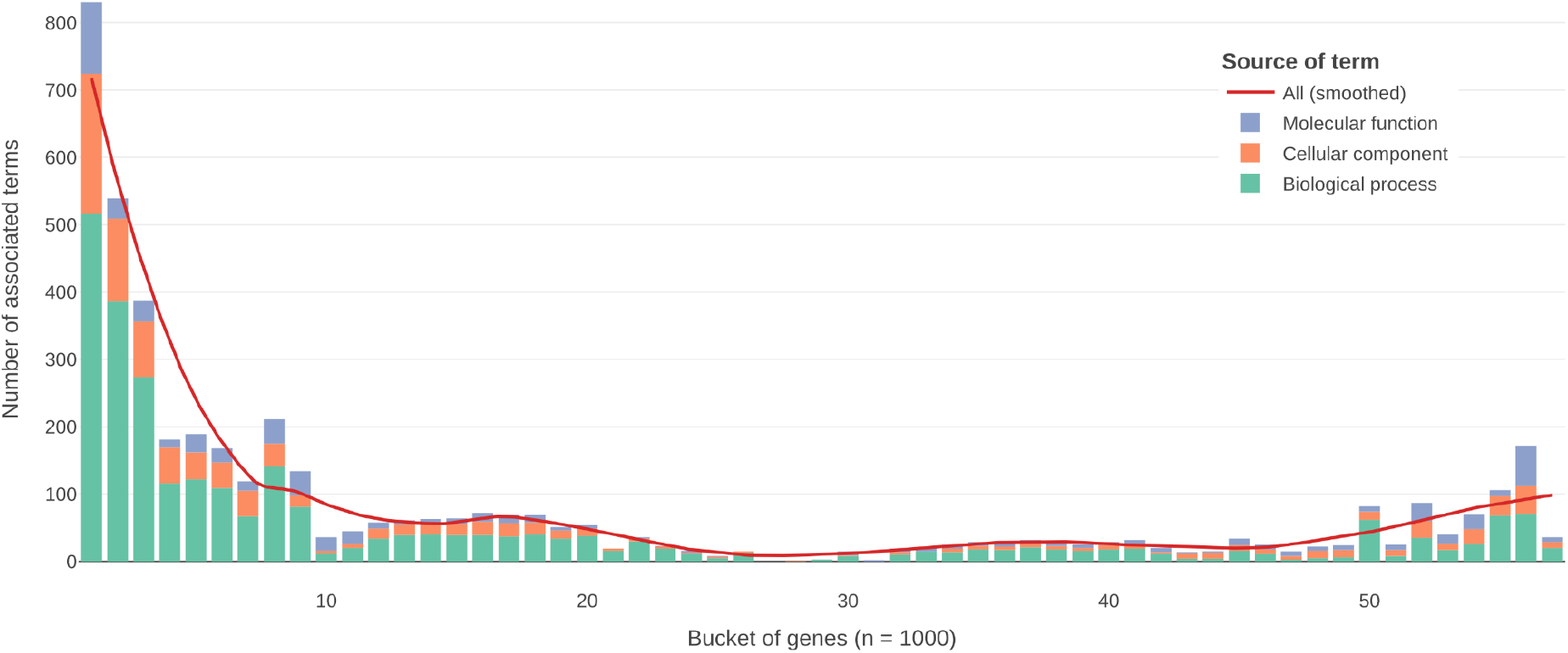
Number of GeneOntology terms for a non-overlapping sliding window of 1000 genes along the ranking of ubiquity. We observe that the most ubiquitous genes have many more known annotations than any other equally-sized window of genes. The genes of average ubiquity have almost no associations with GeneOntology terms. The number of associations rises again for the least ubiquitous genes. The “Additional information” section on the website https://ubigen.uni-rostock.de includes an extended interactive plot that uses a smaller bucket size and shows all available term sources from g:Profiler.

In some studies, the terms housekeeping or ubiquity are used in the context of a limited coverage that only considers a specific set of tissues and/or conditions (like, e.g., cancer^10^ or Parkinson’s disease^11^), to describe genes that are characteristic for that situation and highly expressed. Depending on the kind of analysis, the selection of ubiquitous genes can be positive as just shown for cancer or negative, when searching for specific explicitly non-ubiquitous marker proteins^12^. Such constraints are not implemented in the current version of the interface.

The ranking conceptually resembles the broader effort of the HRT atlas^13^ and the recent in depth analysis of ubiquitous genes^2^. Our approach contributes an interactive user interface and the extra confidence from Figure 2 with details provided online that these genes, and the derived ranking for their ubiquity, are indeed meaningful for both health and disease – on both ends of the spectrum.

## Acknowledgements

The authors thank Georg Fuellen for comments to the manuscript.

